# Identification and quantification of α- and β-amanitin in wild mushrooms by HPLC-UV-EC and HPLC-DAD-MS detection

**DOI:** 10.1101/2022.03.09.483521

**Authors:** Isabel Barbosa, Cátia Domingues, Fernando Ramos, Rui M. Barbosa

**Author notes:** Corresponding author: Isabel Barbosa, *E-mail address.

## Abstract

Amatoxins are a group of highly toxic peptides, which include α- and β-amanitin, found in several species of mushrooms (e.g. *Amanita phalloides*). Due to their high hepatotoxicity, they account for most deaths occurring after mushrooms ingestion. The determination of α- and β- amanitin content in wild mushrooms is invaluable for treating cases involving poisoning. In the present study, we have developed and validated an analytical method based on high-performance liquid chromatography, with in-line ultraviolet and electrochemical detection (HPLC-UV-EC), for the rapid quantification of α- and β-amanitin in wild mushroom samples collected from the Inner Center of Portugal. A reproducible and simple solid-phase extraction (SPE) using OASIS^®^ PRIME HLB cartridges was used for sample pre-treatment, followed by chromatographic separation based on the RP-C18 column. The UV and EC chromatograms of α- and β-amanitin were recorded at 305 nm and +0.600 V *vs*. Ag/AgCl, respectively. The linear quantification for both amanitins was in the range of 0.5–20.0 μg·mL^-1^ (R^2^ > 0.999). The LOD, calculated based on the calibration curve, was similar for UV and EC detection (0.12-0.33 μg ml.^-1^). Intra-day and inter-day precision were less than 13%, and the recovery ratios ranged from 89% to 117%. Nine *Amanita species* and five edible mushrooms were analysed by HPLC-UV-EC, and HPLC-DAD-MS confirmed the identification of amatoxins. We find high α- and β-amanitin content in *A. phalloides* and not in the other species analysed. In sum, the developed and validated method provides a simple and fast analysis of α- and β-amanitins contents in wild mushrooms and is suitable for screening and routine assessment of mushroom intoxication.

**Highlights:**

- New validated method using HPLC-UV-EC to determine α- and β-amanitin in wild mushrooms.
- Reproducible and fast SPE procedure for small samples.
- Effective sample pre-treatment with the OASIS^®^ PRIME HLB SPE cartridge.
- Identification and quantification of α- and β-amanitin in wild mushroom samples from Portugal.
- HPLC-DAD-MS confirmation of amatoxins present in mushroom samples.

**Graphical abstract:** 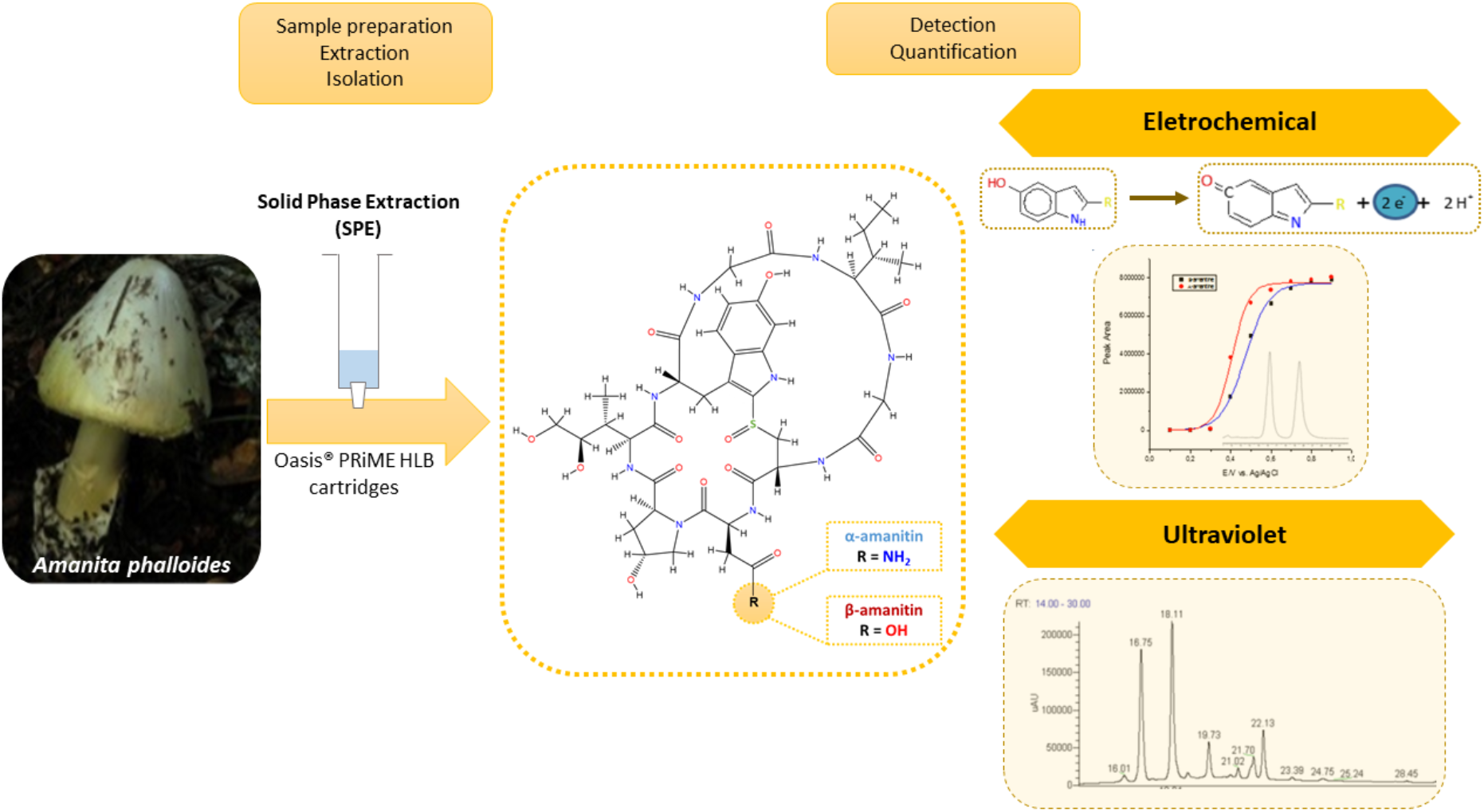

## 1. Introduction

Mushrooms are of great interest in food and pharma industries due to their well-known nutritional (1) and medicinal properties (2–6). Mushrooms have also been used on mycoremediation as they can degrade different types of pollutants, being considered “*clean technologies*” (7). Although, ingestion of some mushroom species by humans is dangerous, causing acute toxicity, which can be life-threatening. Poisoning by mushrooms (micetism) containing amatoxins accounts for over 90% of the fatalities (8), and the incidence is probably underestimated due to a large number of unreported cases (9). The severity of mushroom poisoning is multifactorial depending on the geographic location, the growth conditions, the amount of toxin ingested, and the genetic profile (10).

Mushroom toxins have been grouped in the following categories: cyclopeptides, orellanine, monomethylhydrazine, disulfiram-like reaction, muscarine, hallucinogenic indoles, isoxazoles, and GI irritants (11). The principal poisonous ingredient of the group of cyclopeptides is the amatoxins, namely the bicyclic octapeptides (Figure 1) from *Amanita species* (11, 12), including α- and β-amanitin, which are 10-20 times more toxic than phallotoxins (13–15). The structural formula of α-amanitin has been elucidated as a cyclic octapeptide whose ring is divided by a sulfoxide bridge from the original cysteine sulfur to the 2-position of the indole nucleus of a tryptophan unit (Figure 1). Besides a-amanitin, a carboxamide, β-amanitin has also been identified and isolated (16–18). An isoleucine side chain in position 6, a trans-4-hydroxyl group at proline in position 2, and a hydroxylated L-isoleucine side chain in position 3 are the three binding sites indispensable for toxic effects of a-amanitin (17).

**Figure 1.**
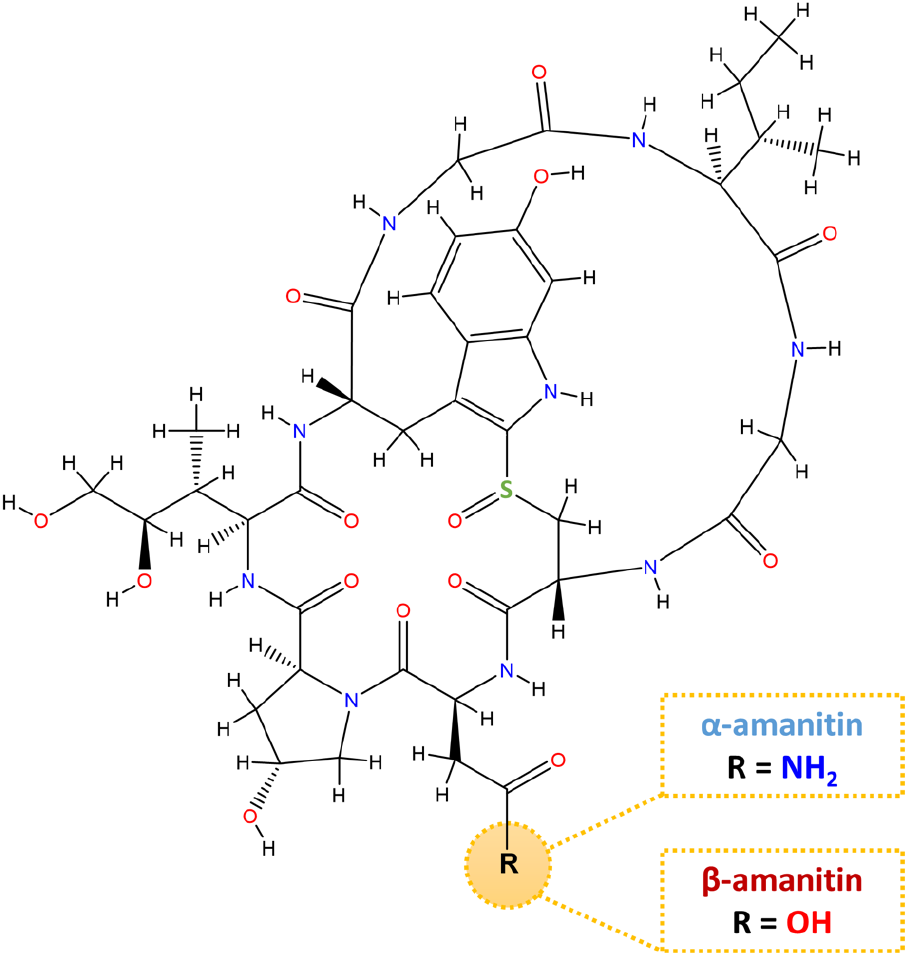
Chemical structure of α- and β-amanitin.

The underlying toxicological mechanisms are usually associated with RNA polymerase II inhibition, which may induce changes in normal cell metabolism, cell death, and tissue necrosis (19). Identifying an antidote remains a challenging task, and innovative therapies are critically needed for the treatment of patients (20–22).

Due to the lethality of amatoxins, and despite the number of methods that have been described, there is still a need for the development of highly sensitive and selective methods that are also simple, fast and accessible for quantifying amatoxins in different matrices such as wild mushrooms or urine and serum.

Several studies have been described for the determination of amatoxins in wild mushrooms based on RP-HPLC with UV detection (23), DAD (24–26), MS-TOF (27), DAD-MS (28–30). To the best of our knowledge, there is no study on the toxin content of various *Amanita species* from the central region of Portugal. Therefore, considering the importance of identification/quantification of these highly toxic amatoxins in mushroom samples, we herein describe a simple, rapid, reproducible, and accurate method, based on RP-HPLC with in-line-UV-EC detection, for identification and quantification of α- and β-amanitin in wild mushrooms collected in the inner centre of Portugal. Moreover, environmental-friendly solid-phase extraction (SPE) using HLB cartridges was firstly used to successfully isolate and quantify α- and β-amanitin from wild mushrooms samples. Nine *Amanita species* and five edible mushrooms were subsequently analysed, and the presence of α- and β-amanitin toxins was confirmed by HPLC-DAD-MS.

## 2. Material and methods

### 2.1. Reagents and Chemicals

All reagents and solvents were of analytical grade unless specified. The standards of α-amanitin (≥90% purity) and β-amanitin (≈90% purity), and sodium acetate were acquired from Sigma–Aldrich (Steinheim, Germany). HPLC-grade methanol and acetonitrile were obtained from Merck (Darmstadt, Germany), and ultrapure water was supplied by a Milli-Q water purification apparatus (Millipore Lda, Bedford, MA, USA). All solutions prepared for HPLC were filtered using a 0.45mm nylon filter. OASIS^®^ PRIME HLB (1mL/30 mg) from Waters Corp. (Mildford, MA, USA) and vacuum manifold system (VacElut 6 Manifold Processing Station, Agilent Technologies, Santa Clara, CA, USA) were used for solid-phase extraction (SPE).

### 2.2. Stock solution for calibration and quality control samples

For calibration and control samples, stock solutions of α- and β-amanitin were prepared by dissolving accurately weighed, amounts of each standard in methanol, obtaining the final concentration of 1.0 mg mL^-1^. Six standard working solutions of α- and β-amanitin (0.5, 1.0, 2.0, 5.0, 10.0 and 20.0 μg mL^-1^) were prepared by diluting the stock solution of each amanitin with the 1 mL of mobile phase [0.1 M sodium acetate, pH 4.7 (A): methanol (B), 83:17, v / v)]. All stock solutions were stored at −20°C in the dark until use. Quality Control (QC) samples were prepared at 0.5 μg mL^-1^ (low-quality control, LQC), 2.0 μg mL^-1^ (medium quality control, MQC), and 10.0 μg mL^-1^ (high-quality control, HQC) for α- and β-amanitin, respectively. The spiked blank shiitake mushroom samples were treated according to the sample preparation described below.

### 2.3. Material of study and sample preparation

#### 2.3.1. Mushroom Collection

Blank shiitake mushroom samples (*Lentinula edodes*) were purchased from local markets. Real samples, fourteen wild mushrooms, were collected in several field trips in the Inner Center Region of Portugal (Figure 2) between September and November. All edible and poisonous mushrooms were properly clean, dried at 40°C for 24 hours, and stored at −20°C.

**Figure 2.**
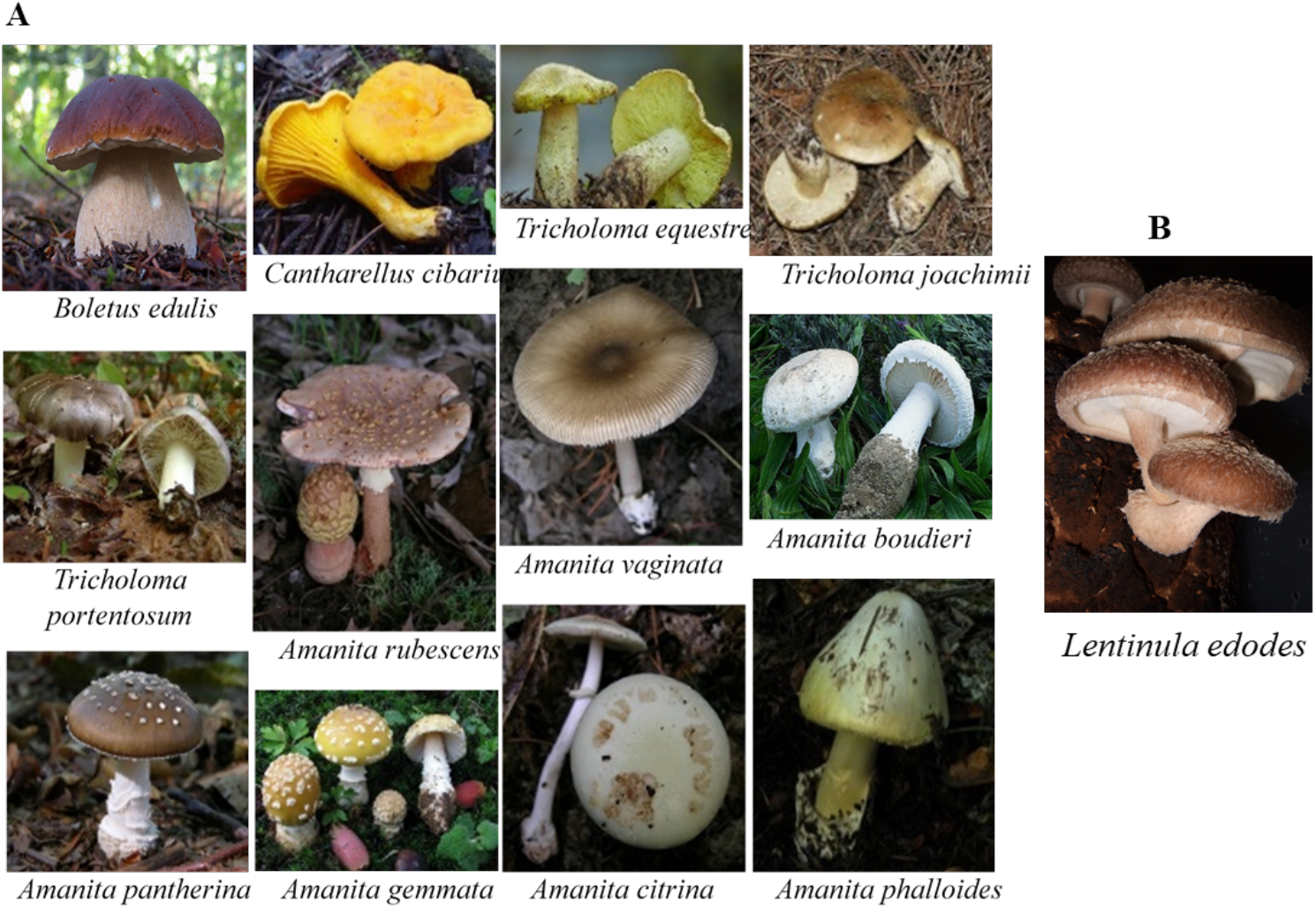
**(A)** Wild mushrooms collected in the Inner Center Region of Portugal and analysed by HPLC-UV-EC and HPLC-DAD-ESI-MS-MS; **(B)** *Lentinula edodes* mushroom used as a blank sample.

#### 2.3.2. Sample Preparation and Extraction Procedure

##### Sample preparation

Sample preparation was performed based on a previously established protocol by Enjalbert et al. (23) with some modifications. Briefly, 0.2 g of all parts of dried mushrooms were dissolved in 10 mL of extraction medium [methanol/water/0.01 M HCl (5:4:1, v/v/v)]. The mixture was vortexed for 30 s, incubated for 1h at room temperature, and centrifuged at 2000 g for 5 min. The supernatant was collected and filtered by a disposable filter holder of 0.45 μm. The filtered solution was evaporated to dryness at 50-55°C and the residue was reconstituted in 10 mL of mobile phase [83 (A); 17 (B)] and then subjected to a solid phase extraction (SPE) procedure carried out on OASIS^®^ PRIME HLB cartridges as shown in Figure 3 and described below.

**Figure 3.**
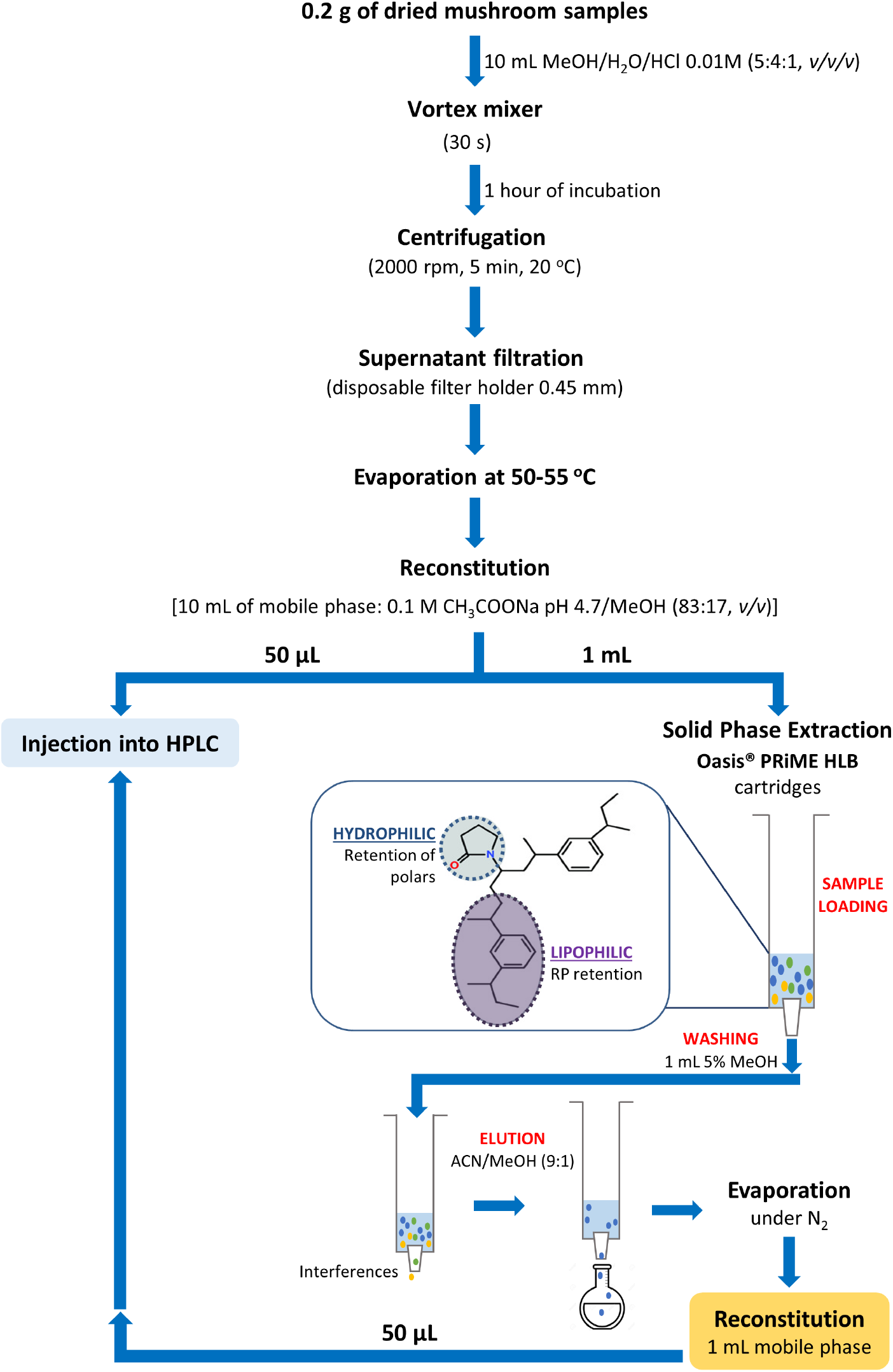
Flow chart showing the proposed sample treatment and analysis for HPLC-UV-EC.

##### Extraction of amanitins

After some optimization steps using different types of cartridges and solvents, the extraction of amanitins was performed using a vacuum manifold system (VacElut 6 Manifold Processing Station, Agilent Technologies, Santa Clara, CA, USA) with a 6 positions rack and OASIS^®^ PRIME HLB cartridges (1 mL/30 mg) were selected. The extraction scheme used to extract the target compounds is illustrated in Figure 3. In brief, 1 mL of the previously obtained solution was loaded onto an OASIS^®^ PRIME HLB extraction cartridge without preconditioning the cartridge sorbent. After, the cartridge was washed with 1 mL of 5% methanol, and the elution was done with 1 mL of acetonitrile/methanol (9:1). The eluate was collected and thoroughly dried to a solid residue under a slight nitrogen flow at room temperature. The residue was reconstituted with 1 mL of mobile phase, and 50 μL was injected into the chromatographic system. Blank shiitake mushroom samples alone and spiked (QC) were used to validate the method, all samples were prepared in triplicated, and the SPE was then performed according to the scheme depicted in Figure 3.

### 2.4. Chromatographic system and conditions

#### HPLC-UV-EC

The chromatographic analyses were carried out using Flexar Perkin Elmer HPLC (Norwalk, USA) chromatography comprising LC Flexar binary pump, LC Flexar autosampler, and LC Flexar detector UV/VIS followed by electrochemical detector BASi (West Lafayette, IN, USA), coupled to Perkin Elmer TotalChrom workstation.

Chromatographic separation was performed at room temperature in a reversed-phase Brisa LC2 C18 column (150 mm x 2.1 mm, 3μm particle size). The mobile phase was composed of a buffer mixture of 0.1 M sodium acetate (pH 4.7) (A) and methanol (B) in a ratio of 83 (A): 17 (B), v / v. Then, it was filtered through a 0.45 μm membrane (Schleicher & Schuell)) and degassed. Isocratic elution was applied at a flow rate of 1.0 mL min^-1,^ and the injection volume was 50 μL for all samples. A sequential detection was performed for all studies using UV detection at 305 nm, followed by EC detection holding potential (+0.600 V *vs*. Ag/AgCl).

#### HPLC-DAD-MS

The analysis of α-amanitin and β-amanitin was carried out on a Surveyor^®^ liquid chromatograph equipped with a photodiode array detector (Surveyor^®^) and interface with a Finnigan LCQ Advantage Ion Max tandem mass spectrometer (Thermo Fisher Scientific, Waltham, MA, USA) equipped with an ESI ionization chamber. Separation was performed on a Spherisorb^®^ ODS-2 C18 reverse-phase column (150 mm x 2.1 mm; particle size 3 μm; Waters^®^ Corporation, Milford, Massachusetts, USA) and a Spherisorb^®^ ODS-2 guard cartridge C18 (10 mm x 4.6 0 mm; particle size 5 μm; Waters^®^ Corporation, Milford, Massachusetts, USA) at 25°C, using 0.1 M sodium acetate pH 4.7 (eluent A) and methanol (eluent B) as mobile phase. The gradient profile was 90% A/10% B (0.0-4.0 min), 82% A/18% B (4.0-10.0 min):0% A/100% B (10.0-30.0 min) and 90% A/10% B (30-40 min) at a flow rate of 200 μL mL^-1^.

The detection was done with a DAD in a wavelength range of 200-600 nm, recorded at 305 nm, followed by a second detection with the hybrid Quadrupole Ion Trap Mass Spectrometer (LCQ Advantage MAX Thermo Finnigan, San Jose, California, USA) operated in the positive electrospray ionization (ESI) mode using selected reaction monitoring (SRM) acquisition. Two consecutive scans were performed: entire mass (m/z 200-1000) and MS^2^ of the most abundant ion in the total mass. Source voltage was 5.00 kV, and capillary temperature and voltage were 150°C and 42V, respectively. Nitrogen was used as nebulizing gas, with a sheath gas flow of 15 (arbitrary unit) and the auxiliary sweeps gas flow of 5 (arbitrary unit). The collision gas was helium with a normalized collision energy of 35%. A precursor ion (MS^1^) and an MS^2^ product ion were obtained for α- and β-amanitin.

### 2.5. Method validation

The HPLC-UV-EC method validation was performed according to US Food and Drug Administration regulations (31), European Medicines Agency (32), and also some complementary aspects taken in the guidelines from the International Conference on Harmonization (33). Validation items include selectivity, linearity, the limit of detection (LOD) and limit of quantification (LOQ), precision (intra- and inter-day repeatability), extraction recovery, and matrix effect.

#### 2.5.1. Selectivity

Selectivity is the ability of the method to assess the analyte(s) unequivocally from interfering components that could potentially exist in the sample. The selectivity of the method was performed by comparing, under optimized chromatographic conditions, chromatograms of standard α- and β-amanitin, extract of blank mushroom samples, and extract of mushroom blank samples spiked with both amanitins to exclude interferences that could co-elute with α- and β-amanitin. The absence of detectable interfering peaks at the α- and β-amanitin retention times was considered as lack of interference.

#### 2.5.2. Linearity

The evaluation of linearity is of great importance. It ensures that in a specific calibration range of concentrations, the instrumental response is proportional to the concentration of the analyte of interest. For this purpose, the linearity response for α- and β-amanitin was obtained by an external calibration using six standards in the mobile phase, injected three times, covering the range 0.5 - 20.0 μg mL^-1^. Linear regression equations evaluated linearity for each curve. Correlation coefficients were obtained, and residual analysis showed the straight-line model is correct (31–33).

#### 2.5.3. Recovery

To ensure that sample extraction is efficient and reproducible, the accuracy of the analytical method was assessed by recovery studies done by spiking the blank matrix with the analyte before sample treatment. Therefore, recovery was used further to evaluate the accuracy of the method in our study and was determined by spiking the shiitake mushrooms (*Lentinula edodes*) with α- and β-amanitin to achieve low (LQC), intermediated (MQC) and high concentrations (HQC), 0.5, 2.0 and 10.0 μg mL^-1^, respectively, by repeated analysis (n=6). Appropriate amounts of working solutions were added to the chopped shiitake mushroom samples for the validation method. After equilibration for 1 h, the validation samples were extracted following the above-mentioned method. The recoveries were determined by comparing the peak areas of spiked samples with that of standard solutions.

#### 2.5.4. Precision

Spiked control quality samples of α- and β-amanitin at three levels of concentration (0.5, 2.0 and 10.0 μg mL^-1^) were tested over the same day to examine the intra-day precision (n=6) of the HPLC system in terms of retention time and peak area. The inter-day precision (n=3) was assessed by testing the same QC samples over three consecutive days. The coefficient of variation (CV) was calculated to estimate precision, according to the following equation: *CV (%) = (σ/μ) x 100*, where σ is the standard deviation and μ is the mean of the response.

#### 2.5.5. Limit of detection (LOD) and limit of quantification (LOQ)

Accordingly, to the standard guidelines for analytical procedures, the LOD is the lowest amount of analyte in a sample that can be detected but not necessarily quantitated as an exact value, and the LOQ corresponds to the lowest amount of analyte in a sample, which can be quantitatively determined with suitable precision and accuracy (31–33). The LOD and LOQ for α- and β-amanitin by the proposed method were determined using calibration standards and calculated using the standard deviation of the residuals (S_y/x_) of the regression line and the slope of the calibration curve (*m*) by the following equations: *LOD* = *3.3(*S_y/x_/*m)* and *LOQ* = *10(*S_y/x_/*m*).

#### 2.5.6. Matrix effects

Matrix effects can dramatically influence analysis performed for identifying and quantifying an analyte (31, 34). The matrix effect was evaluated by comparing the calibration graphs by spiking shiitake mushroom samples with known amounts of α- and β-amanitin, and the calibration graph obtained from α- and β-amanitin standard solutions at the same concentration, by the following equation: matrix effect *(%) = (m_QC_/m_ST_) x 100*, where *m_QC_* is the slope of fortified shiitake mushroom samples, and *m_ST_* is the slope of standard solutions of α- and β-amanitin.

### 2.6. Method application

The developed and validated method was applied to fourteen species of mushrooms collected in the Central region of Portugal, some of them shown in Figure 2. According to the method, all samples were pre-treated, extracted by SPE, and 50 μL of the residue was injected into HPLC-UV-EC to confirm and quantify the presence of these toxins. The identification of the α- and β-amanitin was made by comparing the retention times (t_R_), UV λ_max_, and MS^n^ data with those of known standard references by HPLC-DAD-MS.

## 3. Results and discussion

### 3.1. Optimization of electrochemical detection

The presence of a 6-hydroxytryptophan, an indole nucleus, in α and β-amanitins, makes it possible to detect these compounds by using an electrochemical detector operating in an oxidative mode (35, 36). The EC detector can be more sensitive and selective than a standard UV detector but requires a more extended stabilization period.

To assess the most suitable electrode holding potential to reduce background current and improve the sensitivity and selectivity, electrochemical studies were conducted using standards solutions of both toxins and a wall-jet cell detector comprising a glassy carbon working electrode at potentials in the range of 0.100 to 1.00 V *vs*. Ag/AgCl reference. Figure 4 depicts the hydrodynamic voltammograms obtained by plotting the peak area versus working electrode potential.

**Figure 4.**
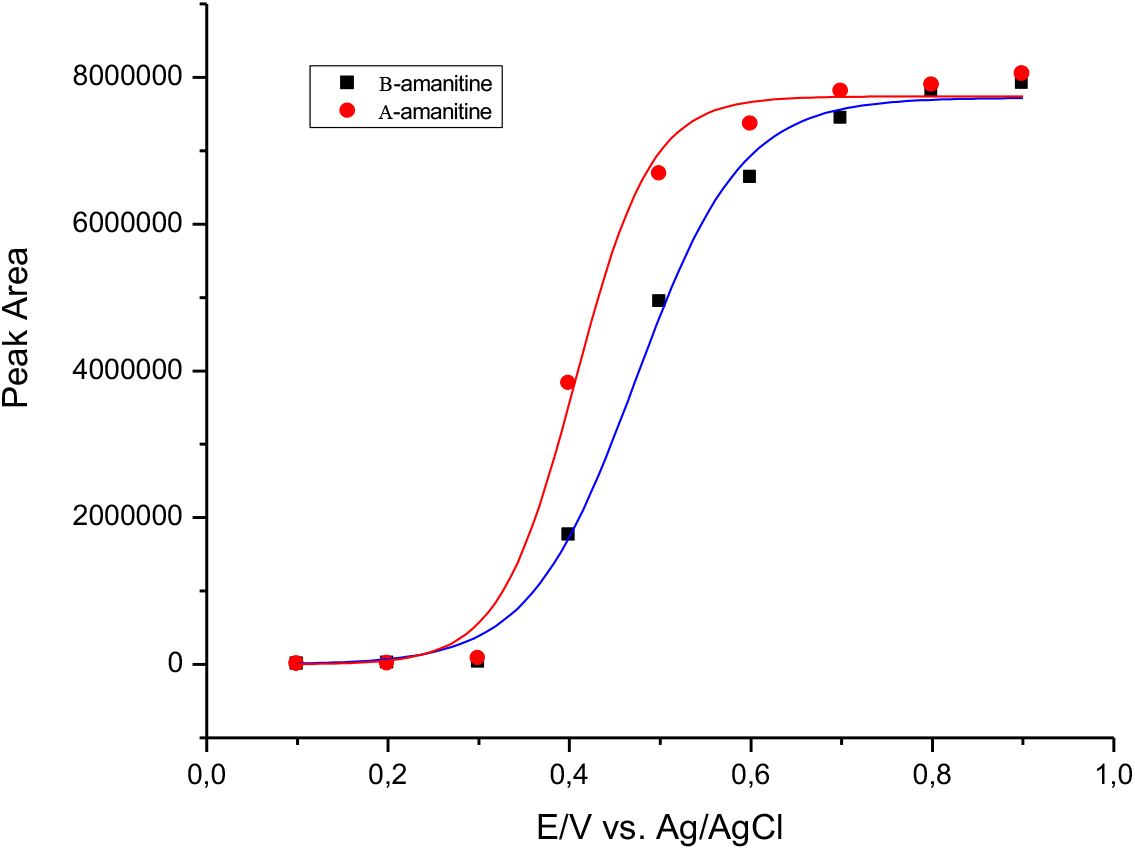
Hydrodynamic voltammograms of α- and β-amanitin standard solutions. Peak areas are plotted *versus* electrode potentials.

The half-wave potentials calculated by fitting data to a sigmoidal function for each curve were very similar for α-amanitin (+ 0.406 V) and β-amanitin (+0.472 V). Therefore, the potential of +0.600 V provides an optimal compromise between sensitivity and selectivity in detecting both amanitins, namely α-amanitin, which is the most toxic molecule. The half-wave potentials are slightly lower (ca. 50 mV) than those reported previously (37). The decrease of the electrode potential to +350 mV is selective and sensitive enough for the determination of α-amanitin in human plasma (Tagliaro et al., 1991). With time the reaction products, mainly derived from the indole residue, tend to adsorb at the working electrode surface leading to a decrease in sensitivity (36). However, this problem can be overcome by regularly polishing the electrode surface. Coulometric detection using working electrodes with high surface areas at oxidation potential of +0.500 V has also been used for the quantification of α-amanitin in urine (35) and in biological samples such as liver and kidney (13).

Figure 5 shows a representative HPLC-EC chromatogram (+0.600 V *vs*. Ag/AgCl) and a chromatogram recorded at 305 nm of α- and β-amanitin with the concentration of 0.5, 1.0 and 2.0 μg mL^-1^ of standard solutions. The retention time was 16.2 s and 19.5 s for β-amanitin and α-amanitin, respectively.

**Figure 5.**
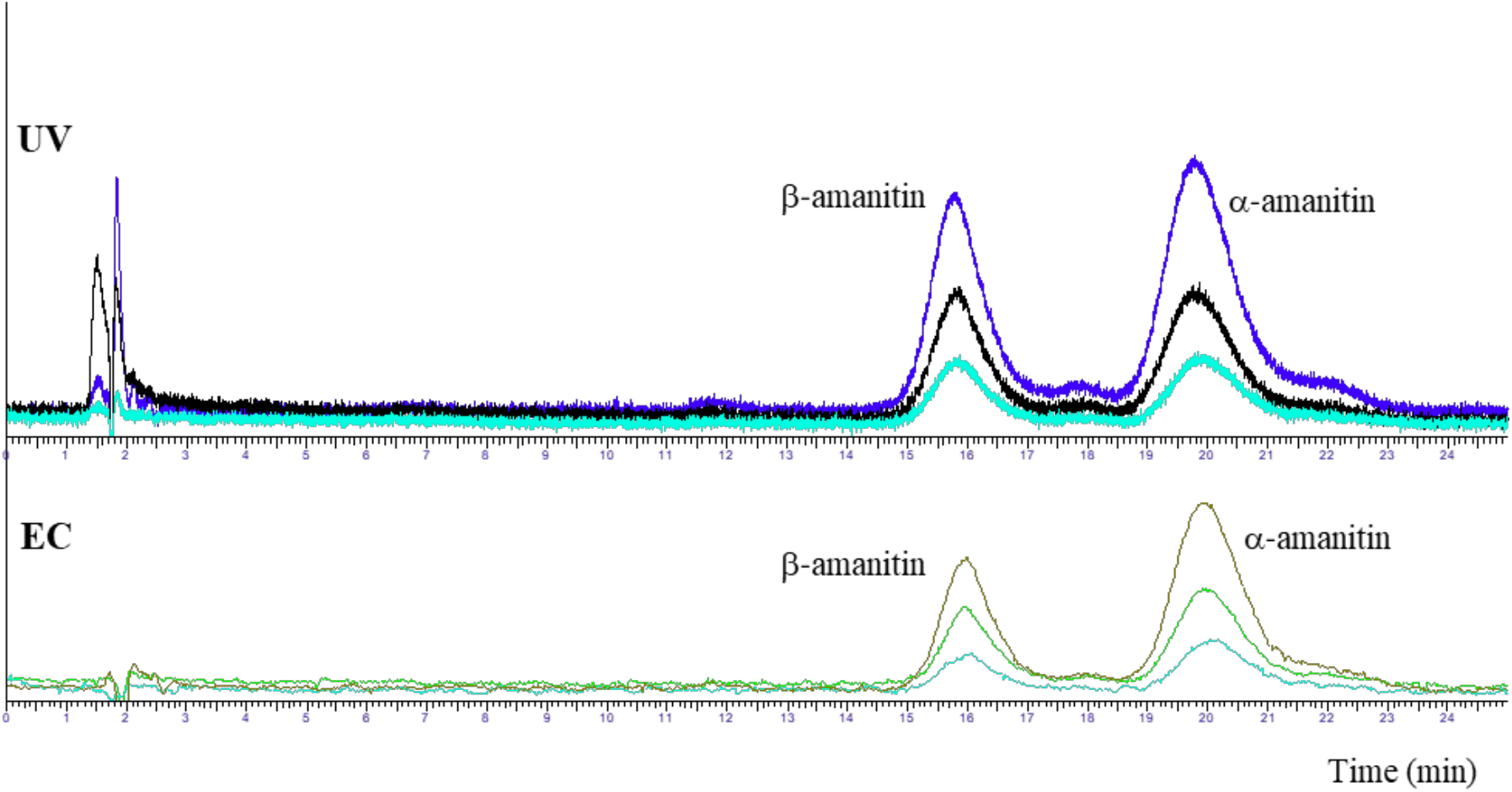
Representative chromatograms of β-amanitin and α-amanitin with the concentration of 0.5, 1.0 and 2.0 μg mL^-1^ of standard solutions recorded by amperometry detection (+0.600 V *vs*. Ag/AgCl) and at 305 nm. The mobile phase composition is a buffer mixture of 0.1 M sodium acetate (pH 4.7) and methanol in a ratio of 83:17 (v/v).

### 3.2. Sample preparation: optimization of extraction and clean-up

The extraction of the analytes of interest from a biological matrix or a mushroom sample is critical in developing an analytical method because it will affect the overall sensitivity and selectivity (38).

Several procedures for the extraction of toxins from mushroom samples have been reported. α-, β-amanitin, and phalloidin have been extracted under acidic conditions such as 0.1% trifluoroacetic acid in methanol (30), 0.5% acetic acid in methanol (39), and 2.5% formic acid in acetonitrile (40), methanol (41), methanol/water (MeOH/H_2_O; 1:1) (42, 43) and acidified methanol/water (MeOH/H_2_O/(HCl or HCOOH) 0.01 M; 5:4:1) by addition of mineral or organic acid (23, 44, 45). Thus, extraction solvents reported in the literature are different according to the mushroom toxins (27).

In the present work, the mushroom sample preparation was performed two steps before the injection into the HPLC system. First, the mushroom samples were crushed and homogenized with acidified aqueous methanolic solutions in accordance with the work of Yoshioka (27). Then, the extracted solution was cleaned up by-one step PRIME (process, robustness, improvements, matrix effects, ease of use). The OASIS^®^ PRIME HLB cartridge is a hydrophilic-lipophilic balance copolymer (HLB) that does not require any solvation, equilibration, or conditioning step, thus saving time and solvent expenses. Being the β-amanitin, an acid compound (-OH group on aspartic residue), elutes first than α-amanitin, a neutral compound (-NH_2_ group on aspartic residue). The recovery rates and sensitivity were higher using methanol. The effective PRIME pass-through clean-up procedure using the Oasis PRIME HLB cartridge was optimized. The results demonstrated higher extraction capacity with recoveries in the range of 80 to 100% (n=5). Literature is scarce on different types of SPE cartridges and purification procedures for cleaning mushrooms complex extracts with medium strength anion exchange (MAX) (40) and HLB copolymer with 40% methanol in water eluent (30, 46).

Our results agree with those reported by Zhang et al. in biological samples (plasma, serum, and urine) for a-amanitin, β-amanitin, g-amanitin, phalloidin and phallacidin (47). The optimized pre-treatment procedure using the OASIS^®^ PRIME HLB method proved to be efficient, provides good recoveries, controls matrix interference, and is time-effective, even without preconditioning. In addition, it is environmentally friendly when compared with other SPE cartridges or other extraction procedures (48).

### 3.3. Optimization of the chromatographic conditions

The chromatographic separation of α- and β-amanitin was carried out by a reverse-phase column Brisa LC2 C18 (150 mm x 2.1 mm, 3 μm) in isocratic conditions. It contains a hydrophobic stationary phase and effectively retains lipophilic analytes, and separates a wide range of compounds (charged/non-charged and polar/apolar). Moreover, it offers an excellent lifetime under extreme pH conditions (including pH 4.7), and it is short length (150 mm), allowing rapid elution of compounds. By recording the absorption spectra of α- and β-amanitin, the wavelength of 305 nm was selected for both compounds allowing high sensitivity and selectivity for both toxins. They contain the 6-hydroxyTrp, which absorb strongly at 305 nm (23, 49). Phosphate or citrate buffers have been used as a mobile phase in separating amanitins (13, 50). Still, they could cause precipitates deposition, creating problems in the HPLC system valves. However, we used acetate buffer (pH 4.7), which in addition to allowing a low background current for EC, eluted β-amanitin first followed by a-amanitin, exhibiting sharp and well-separated peaks, at a flow rate of 1 mL min^-1^ in an overall analysis time of 25 min. The chromatographic conditions used for HPLC-DAD-MS were also optimized to obtain an efficient separation and identification of amanitins in wild mushroom samples. In addition, several MS parameters have been optimized for greater sensitivity.

### 3.4. Validation

#### 3.4.1. Selectivity/specificity

Under optimized conditions, the retention time of β- and α–amanitin were ca. 16 min and 19 min, respectively. Blank shiitake mushroom samples, alone and spiked with α- and β-amanitin, presented no interfering peaks at the α- and β-amanitin retention times, indicating the excellent selectivity of the method.

#### 3.4.2. Linearity

The calibration curves, by using standard solutions with six different concentrations of each amanitin, were linear over the range of 0.5 to 20.0 μg mL^-1^, with correlation coefficients (R^2^) greater than 0.9990 (Table 1).

**Table 1.**
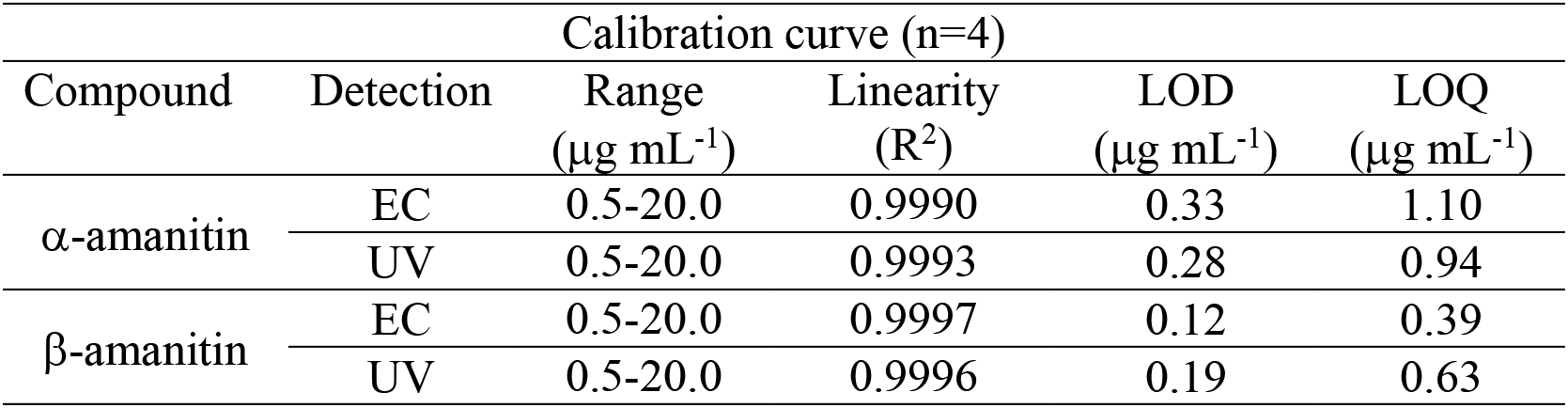
Limits of detection (LOD), limits of quantification (LOQ), and calibration parameters for α- and β-amanitin by HPLC-UV-EC.

| Compound | Detection | Calibration curve (n=4) |  |  |  |
| --- | --- | --- | --- | --- | --- |
| | | Range<br>($\mu\text{g mL}^{-1}$) | Linearity<br>($R^2$) | LOD<br>($\mu\text{g mL}^{-1}$) | LOQ<br>($\mu\text{g mL}^{-1}$) |
| α-amanitin | EC | 0.5-20.0 | 0.9990 | 0.33 | 1.10 |
|  | UV | 0.5-20.0 | 0.9993 | 0.28 | 0.94 |
| β-amanitin | EC | 0.5-20.0 | 0.9997 | 0.12 | 0.39 |
|  | UV | 0.5-20.0 | 0.9996 | 0.19 | 0.63 |

#### 3.4.3. LOD and LOQ

The LOD and LOQ were evaluated, and the data are shown in Table 1. Based on the calibration curve, the LOD and LOQ values of α-amanitin were slightly higher than those of β-amanitin. The comparison between the two detectors (UV and EC) shows no significant difference in the determination of that LOD and LOQ for α- and β-amanitin. However, electrochemical detection complements the method, improving selectivity compared to UV detection in the wavelength of 305 nm selected for both amanitins.

According to the literature, the analytical methods used for the analysis of α- and β-amanitin, especially α-amanitin, based on HPLC with electrochemical detection are scarce and all based on different and biological samples (35–37). It should be noted that comparing our LOD/LOQ values with those reported in the literature is challenging, because methodological details are often poorly described or omitted in most of them. Moreover, it is important to emphasize that the current work is focused on mushroom samples and not on biological samples, and poor information was recorded.

The limits of detection (LOD) obtained for α- and β-amanitin and those described in the literature for mushroom extracts are shown in Table 2.

**Table 2.** Limits of detection for direct mushroom extracts.

| Matrix | Mushroom toxin | LOD (ng g<sup>-1</sup>) | Method | Reference |
| --- | --- | --- | --- | --- |
| Mushroom tissue | α-amanitin<br>β-amanitin | EC-640 ng g<sup>-1</sup><br>UV-207 ng g<sup>-1</sup><br>EC-380 ng g<sup>-1</sup><br>UV-940 ng g<sup>-1</sup> | HPLC-UV-EC | Present Work |
| Mushroom tissue | α-amanitin<br>β-amanitin | 20 ng g<sup>-1</sup><br>20 ng g<sup>-1</sup> | HPLC-ESI-MS | (40) |
| Mushroom tissue | α-amanitin<br>β-amanitin | 30 ng g<sup>-1</sup><br>30 ng g<sup>-1</sup> | HPLC-TOF-MS | (30) |
| Mushroom tissue | α-amanitin<br>β-amanitin | 230 ng g<sup>-1</sup><br>190 ng g<sup>-1</sup> | HPLC-TOF-MS | (27) |
| Mushroom tissue | α-amanitin<br>β-amanitin | 2 ng g<sup>-1</sup><br>2 ng g<sup>-1</sup> | HPLC-TOF-MS | (51) |

As can be observed in Table 2, HPLC-DAD-MS showed LODs in the range of 2-230 ng g^-1^ based on a signal-to-noise of 3. It should be noted that the LOD and LOQ obtained in the present work were calculated based on the calibration curve and produced more conservative estimates.

#### 3.4.4. Recovery and matrix effect

Table 3 summarizes the overall recovery and the matrix effect of HPLC-UV-EC assay of α- and β-amanitin for the shiitake mushroom extracts, expressed as the percentage of recovery and relative standard deviation (%RSD). As shown, recoveries of a-amanitin ranged from 93% to 117% and 89% to 114% for β-amanitin, with an RSD < 5% for both amanitins. The method presents a good extraction efficiency with reproducible recovery since the losses of both amanitins were minimal during the extraction procedure, even for the lowest concentration.

**Table 3.** Recovery and matrix effect values for α- and β-amanitin analysed by HPLC-UV-EC.

| Compound | Detection | Concentration<br>($\mu\text{g mL}^{-1}$) | Recovery<br>(%) (RSD) | Matrix effect<br>(%) (RSD) |
| --- | --- | --- | --- | --- |
| α-amanitin | EC | 0.5 | 117 (4.78) | 96.9 (8.01) |
|  |  | 2.0 | 99 (1.37) |  |
|  |  | 10.0 | 96 (1.48) |  |
|  | UV | 0.5 | 104 (3.72) | 95.8(8.9) |
|  |  | 2.0 | 93 (1.64) |  |
|  |  | 10.0 | 97 (0.34) |  |
| β-amanitin | EC | 0.5 | 114 (3.25) | 95.8(11.4) |
|  |  | 2.0 | 103 (2.94) |  |
|  |  | 10.0 | 94 (0.48) |  |
|  | UV | 0.5 | 92 (3.9) | 97.6 (10.9) |
|  |  | 2.0 | 89 (2.7) |  |
|  |  | 10.0 | 93 (0.6) |  |

The matrix effect was evaluated by comparing the slopes of calibrations curves obtained when processing spiked shiitake mushroom samples and standard solutions of α- and β-amanitin at the same concentrations. The results obtained were in the range from 95.8% to 97.6% for low, medium, and high concentrations of α-and β-amanitin, respectively (Table 3) with RSD <9% for α-amanitin and RSD <12% for β-amanitin. As shown, the results of extraction recovery and matrix effects demonstrated good extraction rates and no significant matrix effects, indicating that the method showed high accuracy.

#### 3.4.5. Precision

As shown in Table 4 the intra-day precision (n=6) for α-amanitin ranged from 0.41% to 5.35% (EC) and from 0.62% to 10.23% (UV), whereas the inter-day (n=3) precision ranged from 7.64% to 12.95% (EC) and from 2.46% to 11.75% (UV). For β-amanitin, the precision of the intra-day ranged from 1.22% to 4.20% (EC) and from 0.69% to 5.33% (UV), whereas the interday precision ranged from 3.81% to 10.38% (EC) and from 3.51% to 5.80% (UV). Acceptable RSD values (<20%) were obtained for all concentrations and both detectors (UV and EC), thus the developed method showed good precision for the determination of α-and β-amanitin.

**Table 4.** Intra and inter-day precision for values for β- and β-amanitin analysed by HPLC-UV-EC.

| Compound | Detection | Concentration<br>($\mu\text{g mL}^{-1}$) | Precision<br>(%RSD) | |
| --- | --- | --- | --- | --- |
|  |  |  | Intra-day | Inter-day |
| α-amanitin | EC | 0.5 | 5.35 | 12.95 |
|  |  | 2.0 | 1.05 | 7.64 |
|  |  | 10.0 | 0.41 | 9.95 |
|  | UV | 0.5 | 10.23 | 3.73 |
|  |  | 2.0 | 1.13 | 2.46 |
|  |  | 10.0 | 0.62 | 11.75 |
| β-amanitin | EC | 0.5 | 4.20 | 10.38 |
|  |  | 2.0 | 3.34 | 5.45 |
|  |  | 10.0 | 1.22 | 3.81 |
|  | UV | 0.5 | 5.33 | 5.80 |
|  |  | 2.0 | 4.55 | 3.69 |
|  |  | 10.0 | 0.69 | 3.51 |

### 3.5. Method application

#### HPLC-UV-EC

In Portugal, between 1999-2008, 93 cases of mushroom poisoning were reported, 63% of which contained amatoxins, leading to 11% of fatal cases (52). The data shows the need to obtain analytical data on the total toxin composition of *Amanita phalloides*, their distribution in the mushroom fruiting body tissues, and their geographic preponderance.

The HPLC-UV-EC analytical method was applied to identify and quantification of α- and β- amanitin in 14 samples of wild mushrooms species. All mushrooms were treated by maceration followed by cleaning with an OASIS^®^ PRIME HLB cartridge, as described in section 2.3, and analysed by HPLC-UV-EC. HPLC-DAD-MS also confirmed the identification of α- and β- amanitin. The analytical method was successfully applied to quantify both amanitins in *A. phalloides* samples.

The results showed the presence of α- and β-amanitin only in the samples of *A. phalloides*. No α- and β-amanitin were detected in any other species analysed, which was confirmed by HPLC-DAD-MS.

The concentration of α- and β-amanitin in the extract of *Amanita phalloides* is shown in Table 5.

**Table 5.**
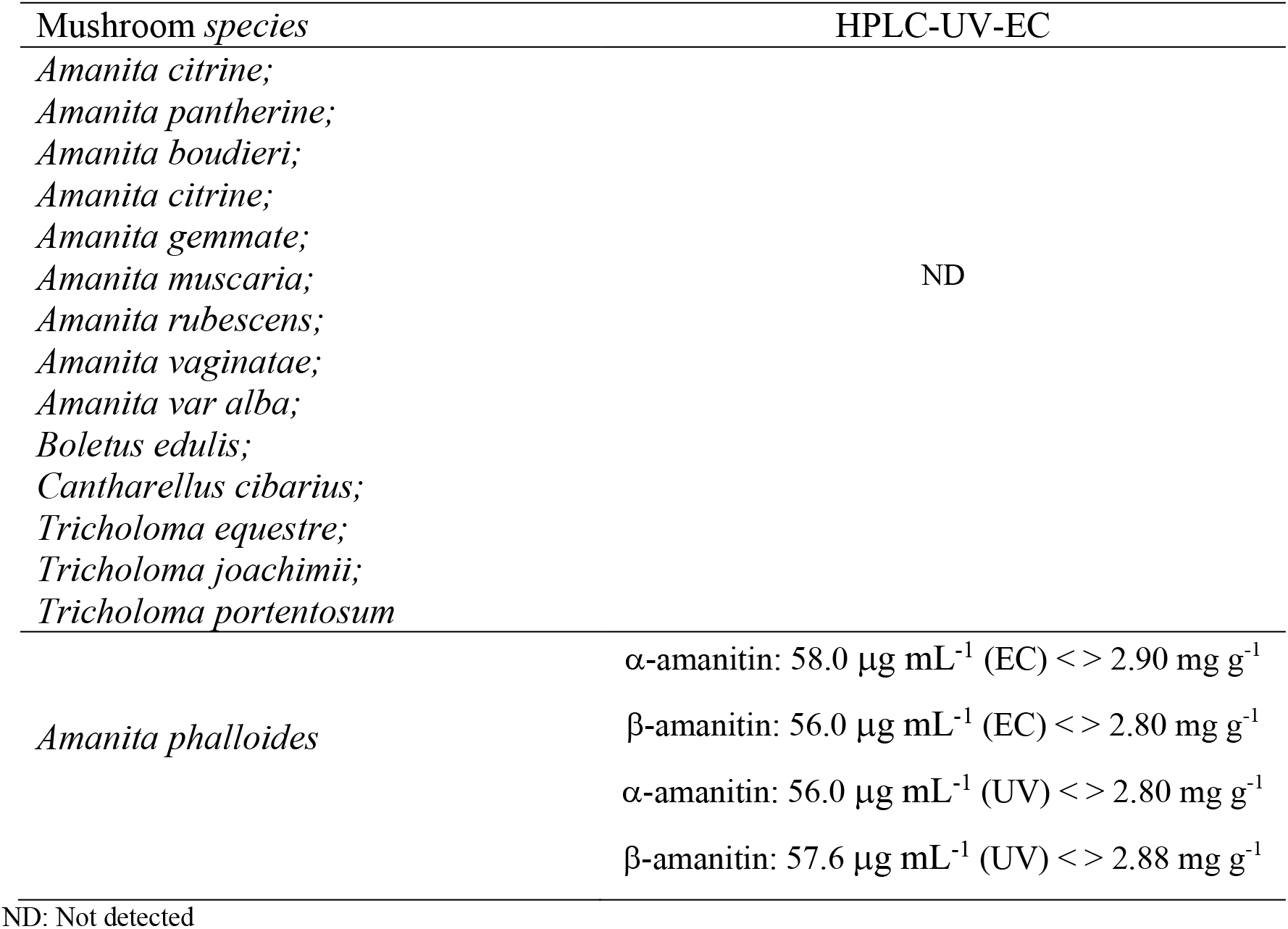
Amanitins identification and quantification in Portuguese wild mushrooms.

As shown in Table 5, the concentration of α- and β-amanitin in *A. Phalloides* was similar for EC and UV detection modes. Our results are in general consistent with previous reports showing that the α- and β-amanitin content of *A. Phalloides* from the Turkye region analysed by HPLC-UV (Kaya 2012, Kaya et. al., 2015). However, the composition and distribution of amatoxins and phallotoxins in different tissues of *Amanita phalloides* from two different locations of Portugal have been already reported by Garcia et al. using LC-DAD-MS. (29). The content and composition of cyclopeptides in lethal amanitins have been shown to vary between different species (43). Previous studies have also shown that climate, topography, soil characteristics and, different fruiting body tissues, and different growth stages influence the concentrations and distributions of amatoxins and phallotoxins in *A. phalloides* (Enjalbert et al. 1996, 1999). The other toxin profiles obtained for the *Amanita species* are derived from the experimental conditions used in the toxin extraction and quantification methods.

#### HPLC-DAD-MS

The chromatogram obtained by HPLC-DAD of the extract of *Amanita phalloides* confirmed the presence of α- and β-amanitin (Figure 6 A). Both compounds exhibited an absorption maximum at 303-305 nm similar to α- and β-amanitin patterns (23). As the backbone of α-amanitin and β-amanitin is a bicyclic octapeptide, and since the only structural difference between α- and β-amanitin resides in the -NH_2_ for a-amanitin and -OH for β-amanitin groups, it is reasonable to expect that both compounds undergo a similar fragmentation pathway under optimized MS conditions. Therefore, we have studied and compared the fragment ions of α- and β-amanitin.

**Figure 6.**
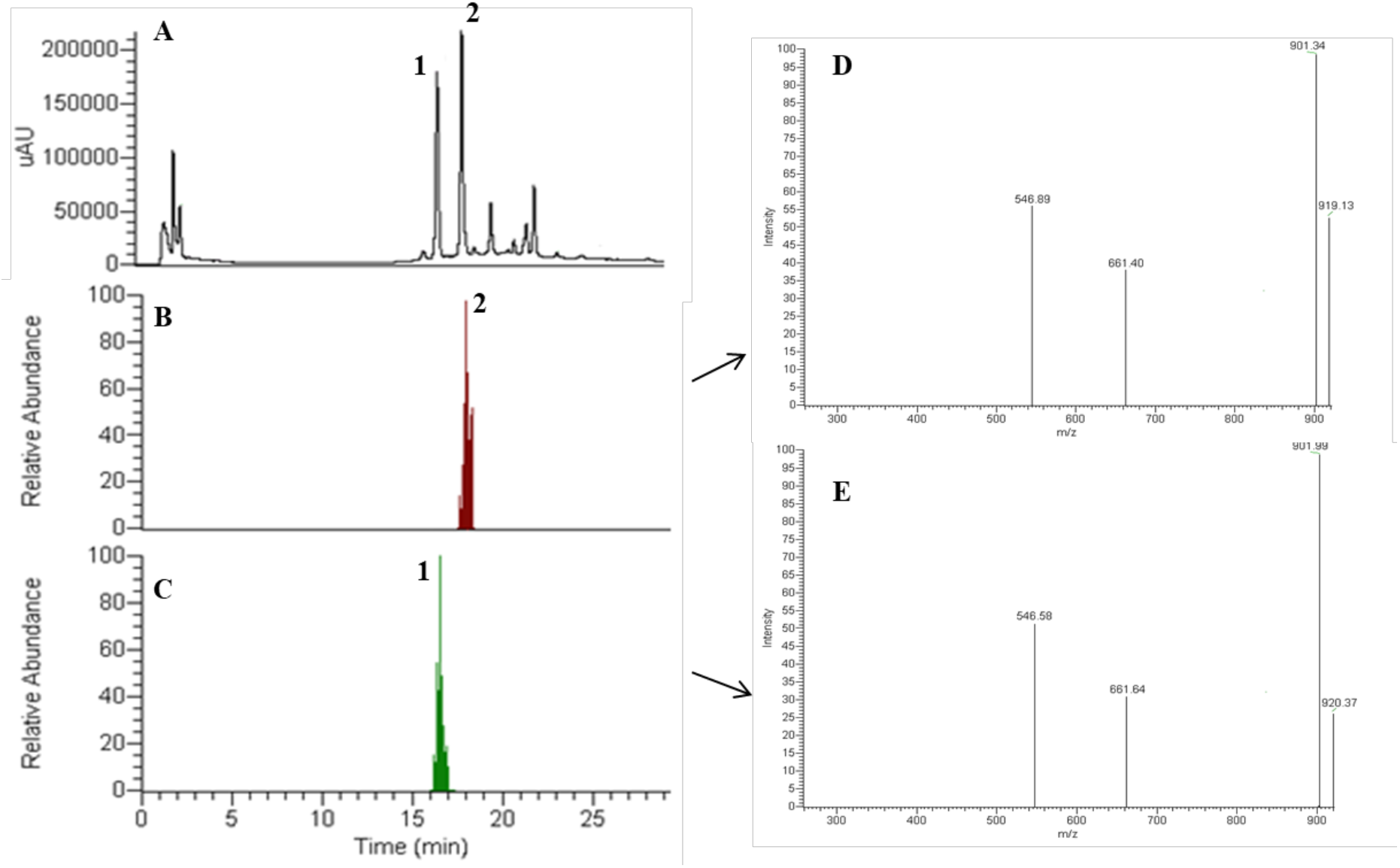
(A) HPLC-DAD chromatogram of an extract of *Amanita phalloides*, monitored at 305 nm, with the separation of peaks: β-amanitin (1) and α-amanitin (2). Total ion chromatogram (TIC) (B) and mass spectrum and fragmentation pattern (D) of peak 2-α-amanitin and total ion chromatogram (TIC) (C) and mass spectrum and fragmentation pattern (E) of peak 1-β-amanitin with a retention time of 18.1 and 16.6 min, respectively.

Based on the chromatographic peaks obtained by DAD and TIC, the molecular ion [M+H]^+^ at m/z 919.00 (relative intensity, 100%) corresponds to the protonated molecular ion of α-amanitin (Figure 6 C). The [M+H]^+^ ion was further selected and isolated to pass through the collision cell to produce MS^2^ fragment ions (Figure 6 E), and the results showed only four fragment ions (Table 6). The limited fragmentation of α- and β-amanitin by the positive ionization mode agrees with a previous work of Johnson et al. (53). The MS^1^ and MS^2^ mass spectra of β-amanitin were acquired under the same conditions, and [M+H]^+^ ion were observed at m/z 920.00 (Figure 6 B, D).

**Table 6.** Retention time and fragment ions of α- and β-amanitin.

| Compounds<br>(Formula) | Molecular<br>weight | Retention<br>time<br>(min) | Protonated<br>Molecular ions<br>(MS<sup>1</sup>, m/z) | Major fragment<br>ions (MS<sup>2</sup>, m/z) |
| --- | --- | --- | --- | --- |
| α-amanitin<br>(C<sub>39</sub>H<sub>54</sub>N<sub>10</sub>O<sub>14</sub>S) | 918.97 | 18.1 | [M+H]<sup>+</sup><br>919.00 | 901.14 (100%)<br>661.43 (44,8%)<br>546.71 (60%) |
| β-amanitin<br>(C<sub>39</sub>H<sub>53</sub>N<sub>9</sub>O<sub>15</sub>S) | 919.95 | 16.6 | [M+H]<sup>+</sup><br>920.35 | 902.09 (100%)<br>661.77 (45%)<br>546.64 (64,8%) |

A comparison of the MS^2^ spectra of α- and β-amanitin fragmented ions revealed some differences, which are summarized in Table 6. The mass spectra for both compounds exhibited base peaks [M+H-H_2_O]^+^ at m/z 901 and 902, respectively, which were produced by dehydration of protonated ions. However, it is not well-known which hydroxyl group is lost from the parent structure when collision-induced dissociation occurs. These fragmented ions are in agreement with previous studies (30, 40, 54–56), and can be used as characteristic ions to identify α- and β-amanitin in complex sample matrices. MS^2^ data also showed fragments at m/z 661 and m/z 547 typical of α- and β-amanitin.

Considering that the same fragment ions were produced, with identical mass differences, it is predictable that α- and β-amanitin have a similar cleavage pathway. In Figure 7 it is proposed a schematic representation of the fragmentation sites in the chemical structure of α-amanitin and the respective characteristic fragments produced (30, 50). Cleavage of peptide bonds at dihydroxy-Ile-Gly (a), Asn-Cys (b) and hydroxyl-Pro-Asn (c) produces the ion peaks at m/z 661 (fragment a to c + H)^+^ fragment at /z 547 (fragment a to b + H)^+^, 373 (fragment b to a + H)^+^, 259 (fragment c to a + H)^+^ and 115 (fragment b to c + H)^+^ which can be observed in spectra of amanitin fragmentation mass.

**Figure 7.**
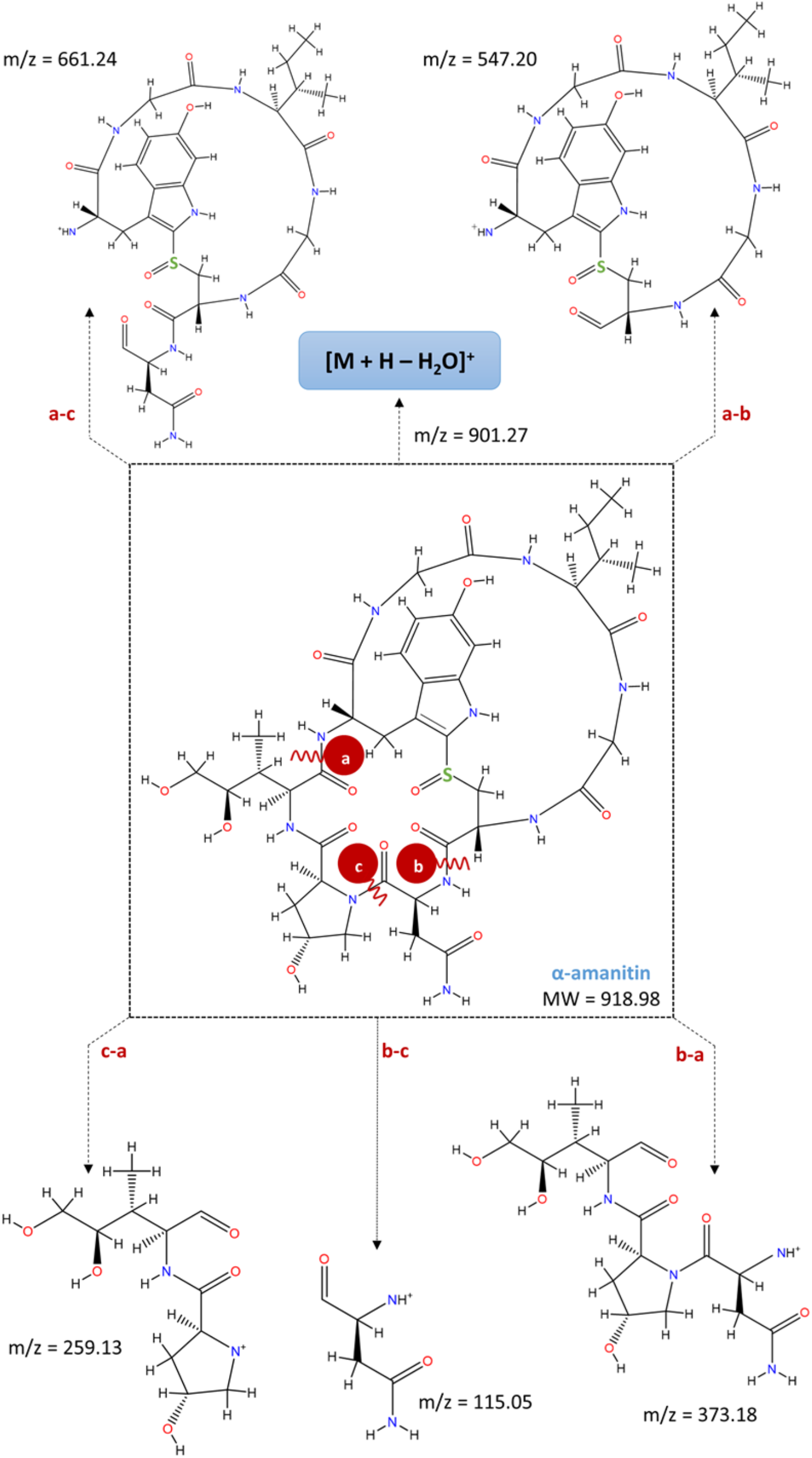
Schematic representation of the fragmentation sites in the chemical structure of α-amanitin and the respective characteristic fragments produced with the corresponding m/z values.

The mass detector in SRM mode for data acquisition provided a high specificity and allowed an unequivocal detection and confirmation of amanitins in wild mushroom samples.

## 4. Conclusions

Developing a simple and accurate analytical method for determining amatoxins content in wild mushrooms is of interest for treating cases involving poisoning. In the present work, we have developed and validated an HPLC method using UV and EC in-line detectors to identify and quantify α- and β-amanitin in samples of Portuguese wild mushrooms. The method proved to be sensitive, selective, accurate, and precise. The proposed method combined with HPLC-DAD-MS successfully confirmed the presence of α- and β-amanitin in the content of *Amanita phalloides* harvested in the central region of Portugal. A fast SPE procedure followed sample preparation by maceration under acidified aqueous methanolic solutions that those not require conditioning or equilibration.

In conclusion, the developed and validated HPLC-UV-EC method can be used in the routine analysis for the detection and quantification of toxins in wild mushrooms. Remarkably, it could be of special interest for the clinical point of view as in circumstances of intoxication by mushrooms such as *A. phalloides*, involving α- and β-amanitin, it may contribute for their identification and quantification, and ultimately, contribute for a specific antidote-therapy execution.

## Acknowledgments

This work was supported by the National Strategic Reference Framework (QREEN) and the European Regional Development Fund (FEDER) through Mais Centro (Programa Operacional Regional do Centro). We are grateful to MSc Marta Leite for her help in collecting and preparing the wild mushroom samples and the manuscript. We also thank MSc Fátima Nunes from the Laboratory of Mass Spectrometry of the University of Coimbra for the technical support in mass spectrometry.

